# Global Efficiency of Retinal Networks shows Robustness and Degenerate States

**DOI:** 10.1101/153742

**Authors:** Camilo Miguel Signorelli

## Abstract

This exploratory project report provides a study of the retina in response to spontaneous and evoked pattern of flashes for different time cycles. These patterns induce changes at individual neural level and network level, while global efficiency, a topological network measure, presents robustness and degenerate states across time. This report also discusses some alternatives to explain changes observed and how it could be related to oscillatory plasticity mechanism.

## Introduction

The retina is a well-structured dynamical neural network. It is considered part of the central system and has interesting anatomically and physiologically properties (Masland, 2012), (Seung & Sümbül, 2014). For example, the retinal network activity can change according to the context (Farrow et al., 2013), inputs are due to rod activity at low luminance (Scotopic condition), or due to cones at high luminance (Photopic condition) (Kolb, 2003; Tikidji-Hamburyan et al., 2015). Furthermore, with or without external stimuli, the retinal network is always presenting neural activity (Evoked or Spontaneous activity). It has two synaptic layers and three nuclear layers (Carcieri et al, 2003; Kolb, 2003; Masland, 2012; Sterling, 2013). Signals from photoreceptor are sent to Bipolar cells (BC) and Horizontal cells (HC), then to Amacrine cells (AC) and Retinal Ganglion cells (RGC) to produce action potentials (Spikes). Retinal cells have a non-lineal signals processing, allowing complex computational capacities (Gollisch & Meister, 2010). For instance, BC can respond differently depending of the presence of ionotropic or metabotropic receptors for glutamate, related with light (ON) or dark (OFF) flashes. Specifically, HC and AC can send different types of signals using various excitatory and inhibitory amino acids, catecholamines, peptides and nitric oxide (Kolb, 2003). These characteristics make of the retina a good model to understand how the brain encodes neural information.

However, what is information for the brain and the way how neural systems codifies information is still an open question and represents a critical problem to understand how the brain works (Breakspear et al, 2010; Gerstner et al, 1997; Gerstner et al, 2007; Quiroga & Panzeri, 2009). Generally speaking, two main types of codification base on spike activity can be mentioned (Cessac et al, 2010; Masuda & Aihara, 2007): i) **Rate codification** (normally spike counts per bin time) and ii) **Temporal codification** (precise time spike latency response). Additionally, independency (Nirenberg et al, 2001) or dependency (Aertsen & Gerstein, 1985; Gollisch & Meister, 2008; Meister, 1996) between neurons can be also assumed.

In general, both spike counts and temporal codification (Schwartz et al, 2007) participate in neural coding. For example, when a repetitive pattern stops (Figure 2), individual RGCs respond strongly depending on the frequency of stimulation (Schwartz & Berry II, 2008) while in the midst of stimulation, the latency firing is constant and the firing rate is smaller than at the end. This type of codification is called Omission Stimuli Response (OSR) (Gao et al, 2009; Schwartz & Berry II, 2008; Schwartz et al., 2007). In other words, the retina recognizes the end of the pattern and the frequency of this pattern. The observed OSR dependency in retina has been attributed to a general resonator neuron mechanism related with sub threshold oscillations (Gao et al., 2009; Schwartz & Berry II, 2008) contrasted with another simple, but not general, explication (Werner et al, 2008). OSR has also been noted in cortical areas of many species, including humans (McAnany & Alexander, 2009); visual system (Bullock et al, 1994; Rogers et al, 1992); auditory system (Klinke et al, 1968) and somato-sensory system (Sutton et al, 1967); whereby OSR could be a mechanism common to nervous system (Bullock et al, 1994; Tiitinen et al, 1994).

These encoding processes have implication on more complex brain areas. For example, the retina has many different types of RGCs which encode different kind of stimuli. It also have a huge spectrum of responses, even for the same stimulus and especially for OSR (Schwartz & Berry II, 2008). In this context, how would other brain areas integrate all this variability? One hypothesis is related with population coding and how the retinal network could keep stable responses to compensate this individual variability.

Dependency and correlations between cells responses may be relevant and related with functional or anatomical connectivity between neurons. In this context, Graph theory, including topological network measures of functional correlation structure, can be applied. Graph theory has been used to study how “healthy” neural network can be characterized (Bullmore & Sporns, 2009; Vincent et al, 2013), in terms of topological measures such as: i) integration, capacity of converge activation patterns, and ii) segregation, capacity of distribute activation patterns across a network (Rubinov & Sporns, 2010). On all these approaches, spiking activity is assumed as the only informative state, even when other types of states could be still informative.

This report will use both independent and dependent coding paradigms to characterize retinal populations responding to repetitive OSR pattern. Thus, it is possible to observe temporal differences on network responses before and after stimulation protocols, mainly changes of latency in ON response, associated with OSR mechanism. It might suggest plasticity effects and dynamical variability of the system. Also, these preliminary results show differences of integration and segregation of neural activity between spontaneous retinal activity and OSR activity. Additionally, interesting conclusions appear after graph analysis of correlations for OSR states, and it shows that topological measures as global efficiency have a robust behavior, producing degenerate states to keep coherence and robustness of the network.

## Methodology

### Animal Model and Recording

Experiments were performed in pieces of retina perfused continuously with Ringer’s AMES medium and obtained from diurnal dichromate rodent ***Octodon degus*** (degu). Degus have ~2.9 million cones with middle wavelength-sensitive (M); 221,000 cones with short-ultraviolet wavelength (UV) sensitive (S), and ~6.5 million rods (Chavez et al, 2003; Jacobs et al, 2003; Palacios & Lee, 2013; Peichl, 2005).

Data is recollected from 3 different animals and 6 pieces of retina in total. Here 429 units are reported and selected from different experiments through visual criteria of spike-triggered average (STA). Experiments were carried under bioethical permit validated by the Universidad de Valparaiso Animal Care Office.

Each retina was dissected and mounted on a 256- multielectrode array (MEA) (Segev et al, 2004) for field and action potentials (spike) recording, using McRack Data Acquisition Software (Multichannel Systems, 2013a, 2013b). The MEA contains 252 recordings, and four ground electrodes arranged in a 16 x 16 layout grid embedded in a transparent glass substrate. The contact to the amplifier is provided by a double ring of contact pads around the rim of the MEA. Electrodes are made of titanium nitride (TiN) with a silicon nitride (SiN) isolator. Contact pads and tracks are made of transparent indium tin oxide (ITO). The average spacing of electrodes was 100 μm in a 16x16 grid. Each electrode has 30 μm diameters with an impedance of approximately 30-50 kΩ. The recording chamber temperature was maintained between 25°C and 30°C.

### Visual Stimulation and Protocols

A Green LED (*λ*_max_ 505 nm, 30 lux) was used for OSR dark or light condition (see below). LEDs were mounted directly underneath the retina and programmed with custom Matlab software.

Experiments consisted of 10 minutes of spontaneous activity recording; 20 minutes of white noise recording; 50 trials of ON-OFF control flashes (Figure 1a), 1 cycle of OSR control for dark or light condition (Figure 1a); 5 cycles of stimulation with 200 trials of green flashes (Habituation protocol) (Figure 1b-e). First four stimulation cycles were separated by break of 2 minutes without simulation activity (it will be called: Resting activity) and the last cycle by break of 4 minutes. Finally, experiments ended with other 50 trials of ON-OFF control flashes.

**Figure 1.**
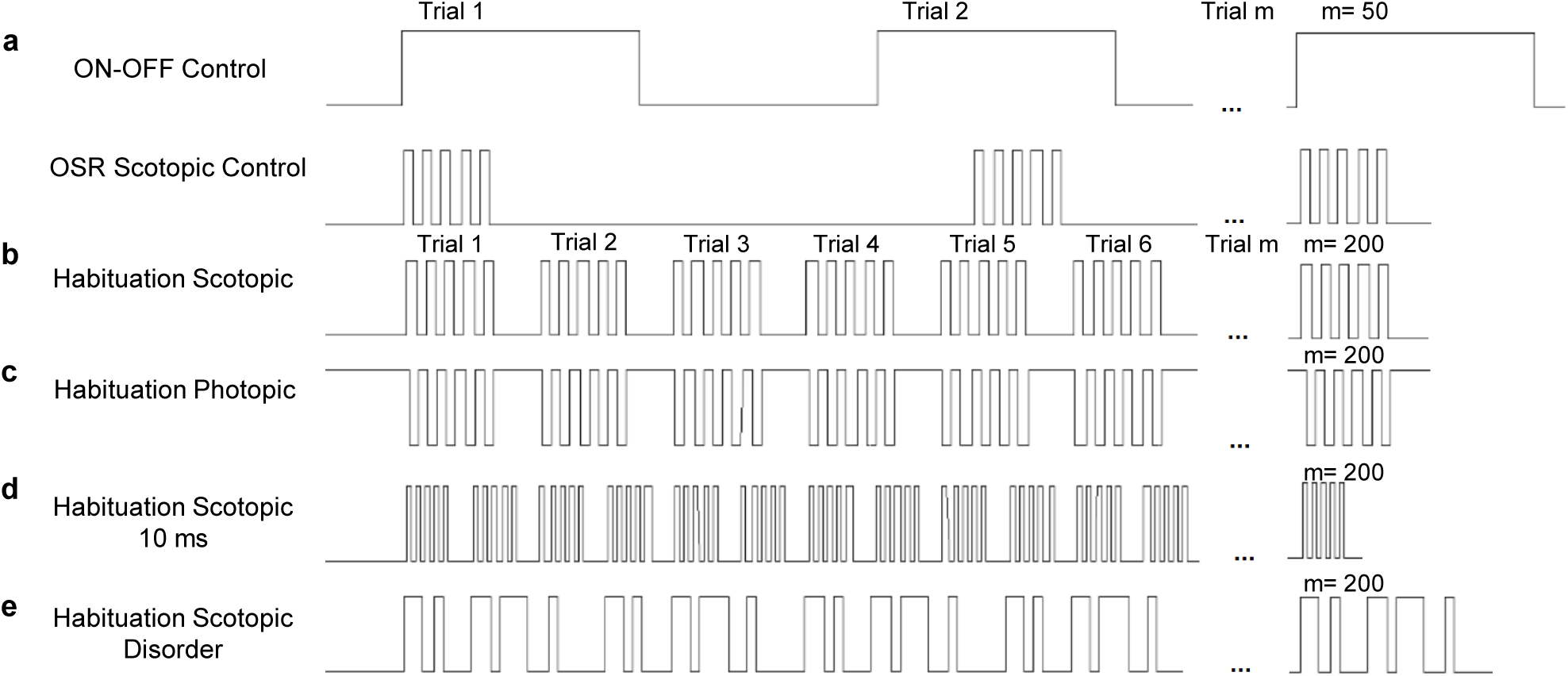
Protocols. a) ON-OFF control pulses had 1 second ON and 1 second OFF. OSR Scotopic (Photopic) Control was with 5 flashes ON of 40 ms and 5 flashes OFF of 40 ms at 12.5 Hz and 2 seconds between trials (for details see Schwartz, G., & Berry II 2008). The number of Trials in both protocols was 50. b) Habituation Scotopic protocol contains 5 ON flashes and 5 OFF flashes, with 40ms duration, i.e. at 12.5 Hz. Each trial was presented at 1.78 Hz, i.e. 160 ms of delay between them and for a total of 560 ms per trial, and a total of 200 trials per cycle with a total of 5 cycles per experiment. First four stimulation cycles were separated by break of 2 minutes without simulation activity (it will be called: Resting activity) and the last cycle by break of 4 minutes. c) Habituation Photopic is similar than Habituation Scotopic, but in Photopic condition. d) Habituation Scotopic 10 ms was a variant with 5 flashes ON of 10 ms and 5 flashes OFF at the same duration. e) Habituation Scotopic Disorder, the same that b) but with different duration of ON and OFF flashes. d) and e) were controls for results of a) and b) experiments. Temporal scale between figures is approximately the same.

The Habituation protocol (Figure 1b-e) contains 5 ON flashes and 5 OFF flashes, with 40ms duration (Schwartz & Berry II, 2008; Schwartz et al, 2007), i.e. at 12.5 Hz. Each trial was presented at 1.78 Hz, i.e. 160 ms of delay between them and for a total of 560 ms per trial, which made each trial dependent, and a total of 200 trials per cycle with a total of 5 cycles (1000 trials in total) per experiment. The cycle for OSR control, dark or light, was similar but only 50 trials and 1 second of delay between them for each condition, which made each trial independent.

Four different variants of Habituation protocol (Figure 1b-e) were tested, and two of them represented controls. i) <u>Habituation scotopic protocol</u> (Figure 1b), where scotopic condition is defined as low luminance before pattern stimulation and resting condition, and light flashes (ON) for stimuli condition (n=3, where n is number of pieces of retina); ii) <u>Habituation photopic</u> (Figure 1c), where photopic is a high luminance condition before pattern stimulation and resting condition, and dark flashes (OFF) in stimuli condition (n=1); iii) <u>Habituation 10 ms</u> (Figure 1d) was a variation of Habituation Scotopic protocol with flashes duration at 10 ms (n=1); iv) <u>Habituation Disorder</u> (Figure 1e) experiment has 5 ON and 5 OFF flashes with different durations between them under scotopic condition, it is not a repetitive pattern within the trial but with a pattern trial to trial (n=1, the protocol was in the same piece that (iv)). Finally, (iii) and (iv) were just controls to make a comparison with the results obtained from the protocols (i) and (ii).

### Sorting, Filters and Classification of units

In order to analyze data, Offline Sorting Software for Plexon Inc was used. A Butterworth 4 poles filter at 100 Hz and threshold of 5 s.d. was used to detect different waveforms. Afterwards, automatic sorting was manually verified (Brown et al, 2004; Lewicki, 1998; Nicolelis, 2008) using a principal component analysis to create clusters depending on type of Waveform (in this case, somatic waveform); Inter Spike Interval Histogram (ISI); autocorrelation and cross correlation between units.

Sorted data was exported to Matlab creating time series (timestamps) for each unit. Verification that RGC were responding at the beginning and at the end of each experimental condition was made. This was to verify the sustained retina response along the experiment.

The spike-triggered average (STA) analysis was run using a custom implementation of the method from (Chichilnisky, 2001). Examples of STA results are shown in Figure S1. Units were removed if their receptive field (RF) showed a peak inferior to 3 s.d. of the background and/or temporal filter curve without ON, OFF or biphasic clear behavior. Two additional filters were applied to verify cell validity and avoid duplication: The first one to calculate correlation between different channels, and other one based in a modulation index calculated as:

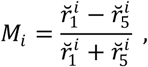

difference between total PSTH firing rates value 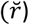 on cycle 1 minus total firing rate value on the final cycle 5 divided by the sum of both values, where *i* is the cell index. If this modulation index was bigger than ± 3 s.d. from the distribution, the unit was discarded.

Finally, each unit was classified through two different criteria: i) ON, OFF or ON-OFF using the PSTH responses over 50 trials of ON-OFF flashes. ON corresponds to an onset response (between 1 ms to 1000 ms and spike count activity >3 s.d. of the total activity); OFF corresponds to the offset response (between 1001 ms to 2000 ms and spike count activity >3 s.d. of the total activity) and ON-OFF has both responses. ii) Using k-means algorithm to classify temporal filter in four main groups: OFF, OFF slow, ON Biphasic, and Biphasic. Finally, the first part of the name in the unit classification corresponds to Pulse (Ps) response and the second part corresponds to the type of temporal filter (Tf). Some examples of responses are in Figure S1.

### Firing Rate and Latency Analyses

A global PSTH is defined as the sum of individual PSTH. It was normalized by the sum of the total activity on five cycles and represented in terms of probability, to make results comparable between different cycles. Adaptation curves are calculated from values of spike counts per trial and the mean corresponds to the whole population.

The mean of firing rate is the mean of activity per unit trial to trial. An OSR peak was calculated with respect to the OSR peak recorded in the first cycle. After that, the time of this peak was kept fixed and it was allowed only a potential variation of ±10 ms during next cycles. The maximal peak was the maximal firing rate value after the last flash, independently of time position. With the goal to measure post OSR activity, it was used the center of mass, defined as the point in the Cartesian coordinate space where the total mass of the system is concentrated. In this case, the center of mass of a post OSR activity was defined as the temporal point after 150 ms after the last flash, i.e. post OSR activity (Pizarro et al, 2014), where the maximal peak of firing rate occurred. The calculus is:

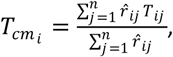

where *i* is the index of cell, *j* is the index trial and *n* is the total number of trials. *T_ij_* is the time-to-peak of the *i* cell in trial *j* and 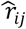 the peak of firing rate after 150 ms. We showed the value 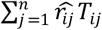 as mass of the center of mass in Figure 3c and 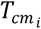 in Figure 3d. The mass is the total counts per bin activity in a temporal range. In addition, to make a simpler comparative analysis and show the percentage of each variation, all these values were normalized according to the first cycle.

To quantify differences on responses between two conditions, initial vs. final Paired Pulse Ratio (PPR) of ON-OFF pulses was calculated by:

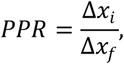

where Δ*x_i_* corresponds to the value at initial condition and Δ*x_f_* is the value at final condition, for one cell. The Δx value was taken directly from the individual PSTH (calculated from 50 trials of ON-OFF control flashes). When Δx was considered the maximal spike counts (peak activity in the temporal window for ON or OFF activity). Finally, three conditions are defined: i) facilitation as: *PPR* > 1, ii) depression as: *PPR* < 1 and iii) unchanged when *PPR* = 1. When Δx was considered time to peak (time to maximal spike count value for ON or OFF activity), facilitation was *PPR* < 1, and depression *PPR* > 1.

### Correlation and Graph Analyses

Timestamps information were converted to binary vectors with zeros and ones corresponding to a bin of 1 ms. The length of these vectors per unit was depending on the temporal window for each analysis (40 ms, 560 ms, 2 minutes etc). Correlation values were computed by a Pairwise Pearson correlation calculated as:

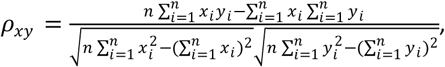

where *i* is *i^th^* component of *x* and *y* vector of *n* length. In other words, Pearson correlation is the covariance divided by the product between standard deviation of both vectors *x* and *y*.

Adjacency matrix was constructed by Pearson correlation (Aertsen & Gerstein, 1985; Vincent et al, 2013) because Maetschke, et. al, 2014 demonstrated that this is one of the best performing simple method of inference in unsupervised networks. The advantage of this method is as accurate as much more complex methods, yet much faster and parameterless (Maetschke, et. al 2014). To calculate the adjacency matrix, length binary vector was taken for every period of interest (Vincent et al., 2013), with 1 ms corresponded to 1 point in the vector. For spontaneous activity condition, 600 seconds are considered. For resting activity and stimulation activity between cycles, a temporal equivalent a 200 repetitions of 560 ms (112 seconds) was used. Then, surrogate data (Grün, 2009; Pipa et al, 2008) was computed based on a randomly dithering of whole spike train against other, because previous works (Grün et al, 2010) showed it as the most robust detector of excess coincidences amongst other surrogates methods. Additionally, jitter length is used with a value of L=50 ms and history length of R=20 ms according to (Harrison & Geman, 2009) and data was shuffled 100 times (number of jitter resample). This surrogate data was used to obtain an absolute threshold value between experimental correlation and random correlation to generate a binary adjacency matrix for each network.

For each cycle, the network was represented by graph characterization, using some measures of integration and segregation. The first one was Global Efficiency (GE) (Latora and Marchiori, 2001; Rubinov & Sporns, 2010).

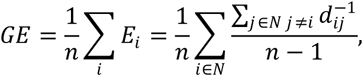

where *E_i_* is the efficiency of node *i*. *N* is the set of nodes which are part of the network and *n* the total number of nodes. *d_ij_* is the shortest path length between nodes *i* and *j* defined for:

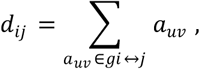

where *g*_*i*↔*j*_ is the shortest path (geodesic) between *i* and *j,* and *a_uv_* is the connection status between *u* and *v*: 1 when the link between *u* and *v* exists (neighbors) or 0 otherwise (also *a_uu_* =0). Finally, note that *d_ij_* = ∞ for all disconnected pairs *i, j*. If *d_ij_* is infinity, GE is zero.

Another measure was Quality of Modularity (Q) (Newman, 2004), defined as:

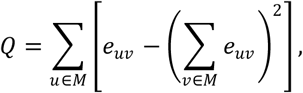

where *u* and *v* are modules within a total of *M* subdivisions of the network and without overlapping. *e_uv_* is the proportion of every link that connects nodes in a module *u* with nodes in module *v*. Modules are functional subgroups of nodes optimally portioned maximizing connection between them. A big value of Q implies more segregation or more subdivision in subgroups than small values of Q.

Finally the variation of information (VI) was computed (Meilă, 2007; Rubinov & Sporns, 2011) between two networks *M* and *M*′, as:

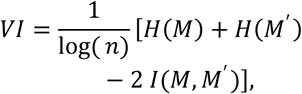

where 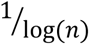 is a scale factor to rescale *VI* in the range [0,1] and *n* the total number of nodes. 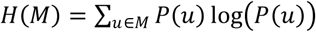 is the entropy definition for the space of partition *M* with 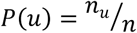 and *n_u_* as the number of nodes in module *u*. The third term is the mutual information defined as:

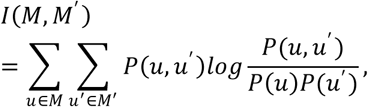

where 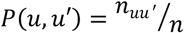 and *n_uu′_* is the number of nodes that are simultaneously in module *u* of partition *M*, and in module *u′* of partition *M*′. This definition implies that *VI* = 0 will mean equal partitions, and *VI* = 1 will correspond to the maximal distance between two partitions (Karrer et al, 2008).

GE, Q, VI and Modules were calculated using Brain Connectivity Toolbox (Rubinov & Sporns, 2010) for Binary Networks.

### Statistical Analyses

Statistic tests were done with Wilcoxon signed rank test for paired data, unless otherwise stated. For effect size analyzes, two different methods were applied depending of the data: i) Mean difference, and ii) Mean difference standardized. The first is defined as:

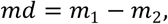

with *m*_1_ the mean of the first distribution, and *m*_2_ the mean of the second distribution. The second method is calculated as:

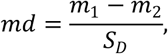

where *S_D_* is the standard deviation of the *difference score* (the differences between matching pairs of data in the groups).

Effect size and Confidence interval was calculated using the effect size Matlab toolbox (Hentschke & Stüttgen, 2011). Confidence intervals of *md* were calculated by a bootstrapping technique (1000 iterations), as a random sampling with replacement (Carpenter & Bithell, 2000; Diciccio & Efron, 1996). Details on the definition of standard error of the mean difference can be seen in (Hentschke & Stüttgen, 2011) and documentation of each method. Finally, to contrast results of the mean distribution shown in Table 1, and avoid problems derived from not normal distribution, It was also applied bootstrapping (3000 iterations) to calculate these values with different distribution assumptions: Normal, Basic, Percentile, and Bias-corrected and accelerated (BCa). This calculation was performed by using R and the Boot function.

**Table 1.** Confidence Interval and mean for different experiments before and after Habituation Protocol.

| Measure | Protocol | Mean Initial | 95% CI Initial | Mean Final | 95% CI Final |
| --- | --- | --- | --- | --- | --- |
| Max Peak OFF | Scotopic | 8.82 | [8.32, 9.32] | 8.83 | [8.28, 9.37] |
| Max Peak ON | Scotopic | 6.8 | [6.2, 7.36] | 7.39 | [6.85, 7.94] |
| Latency Peak OFF | Scotopic | 77.62 | [74.03, 81.22] | 73.26 | [70.01, 76.52] |
| Latency Peak ON | Scotopic | 62.27 | [58.85, 65.7] | 68.92 | [65.42, 72.42] |
| Max Peak OFF | Photopic | 5.52 | [4.8, 6.23] | 5.45 | [4.64, 6.26] |
| Max Peak ON | Photopic | 8.65 | [7.95, 9.36] | 8 | [7.21, 8.78] |
| Latency Peak OFF | Photopic | 71.79 | [60.16, 83.42] | 69.13 | [57.32, 80.94] |
| Latency Peak ON | Photopic | 88.52 | [80.33, 96.71] | 95.81 | [90.77, 100.86] |
| Max Peak OFF | Scotopic 10 ms | 9.98 | [8.75, 11.03] | 9.61 | [8.52, 10.70] |
| Max Peak ON | Scotopic 10 ms | 7.08 | [6.07, 8.88] | 8.38 | [7.29, 9.47] |
| Latency Peak OFF | Scotopic 10 ms | 107.89 | [98.83, 116.96] | 103.25 | [93.73, 111.03] |
| Latency Peak ON | Scotopic 10 ms | 68.17 | [60.15, 76.20] | 75.89 | [67.78, 112.77] |
| Max Peak OFF | Scotopic Disorder | 9.61 | [8.52, 10.70] | 9.51 | [8.50, 10.51] |
| Max Peak ON | Scotopic Disorder | 8.38 | [7.29, 9.47] | 9.12 | [7.82, 10.42] |
| Latency Peak OFF | Scotopic Disorder | 103.25 | [93.73, 112.77] | 99.69 | [90.28, 109.09] |
| Latency Peak ON | Scotopic Disorder | 75.89 | [67.78, 84] | 71.43 | [65.19, 77.67] |

## Results

This work has carried two main types of analyses and results: (i) spike counts and time to peak analyses of groups and individual neurons, which assumes independence on individual neuron responses, and (ii) Graph and Correlation analysis, assuming dependence on neuron responses.

### Characterization of responses for Habituation Protocol

Habituation protocol corresponds to a pattern of flashes which is shown at some frequency, in this case 12.5 Hz, with only 160 ms of delay between trials inducing a second frequency order of 1.78 Hz (see methods Habituation protocol). Two hundred trials were presented in five different time cycles as a lineal sequence. We intersperse activity cycles between cycle 1 and cycle 4 by 2 minutes without activity (Resting activity), and cycle 4 and cycle 5 by 4 minutes. Thanks to this protocol, it was expected to generate habituation without inducing fatigue (see methods, sorting and filters).

Preliminary results showed as some units changed (Figure 2b, c) their activity between cycle 1 and cycle 5, especially on temporal responses (red lines). While others units do not present strong time differences but present some decrease in the spike counts (Figure 2a). Taking 312 ON-OFF units from 4 pieces of retina, it is possible to characterize the Group dynamic of firing rate activity trial to trial between cycle 1 and 5. Adaptation curves trial to trial (Figure 3a) in cycle 1 present a faster decrease at the beginning and then a stabilization. In the next cycles adaptation curves are increasing cycle to cycle, as temporal decay constant shows, until the last cycle, where there is a decrease in the value (Figure 3a). In other words, this protocol is apparently inducing dynamical modifications cycle to cycle.

**Figure 2.**
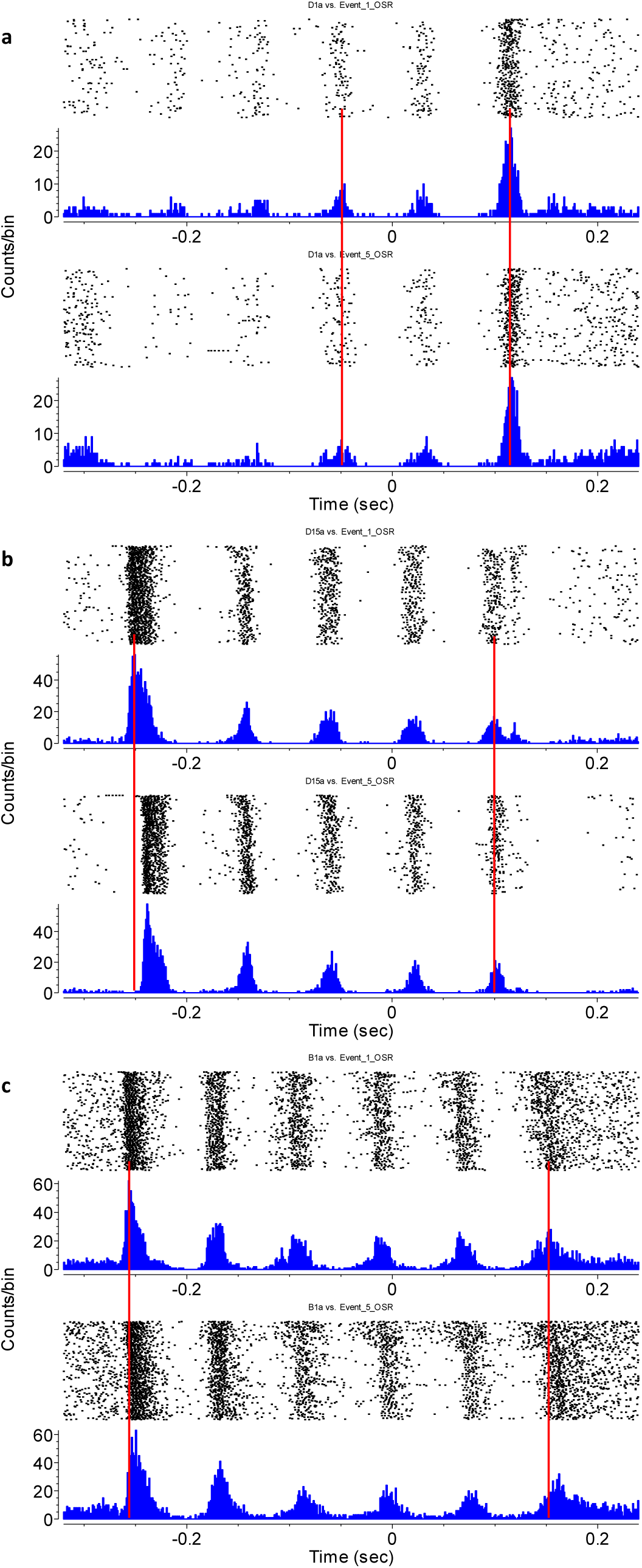
Individual comparison between responses to cycle 1 to cycle 5 in Habituation Protocol. Analyses with individual PSTH show changes between experimental cycles. These changes are principally on time response of some firing rate peaks. a) Unit type OFF with a qualitative decrease on the amplitude of firing probability during flashes but no change on OSR peak, neither latency of its peak. b) Unit type ON with changes on latency response of the first flash. c) Unit type ON-OFF with a shift on latency response for every flash. Units in a-b) were from Scotopic Habituation Protocol and Unit c) from Photopic Protocol. The zero point marks the beginning of last flash and red lines show variation between upper PSTH (Cycle 1) and below PSTH (Cycle 5). Each line in raster plot corresponds to one trial from up to down.

**Figure 3.**
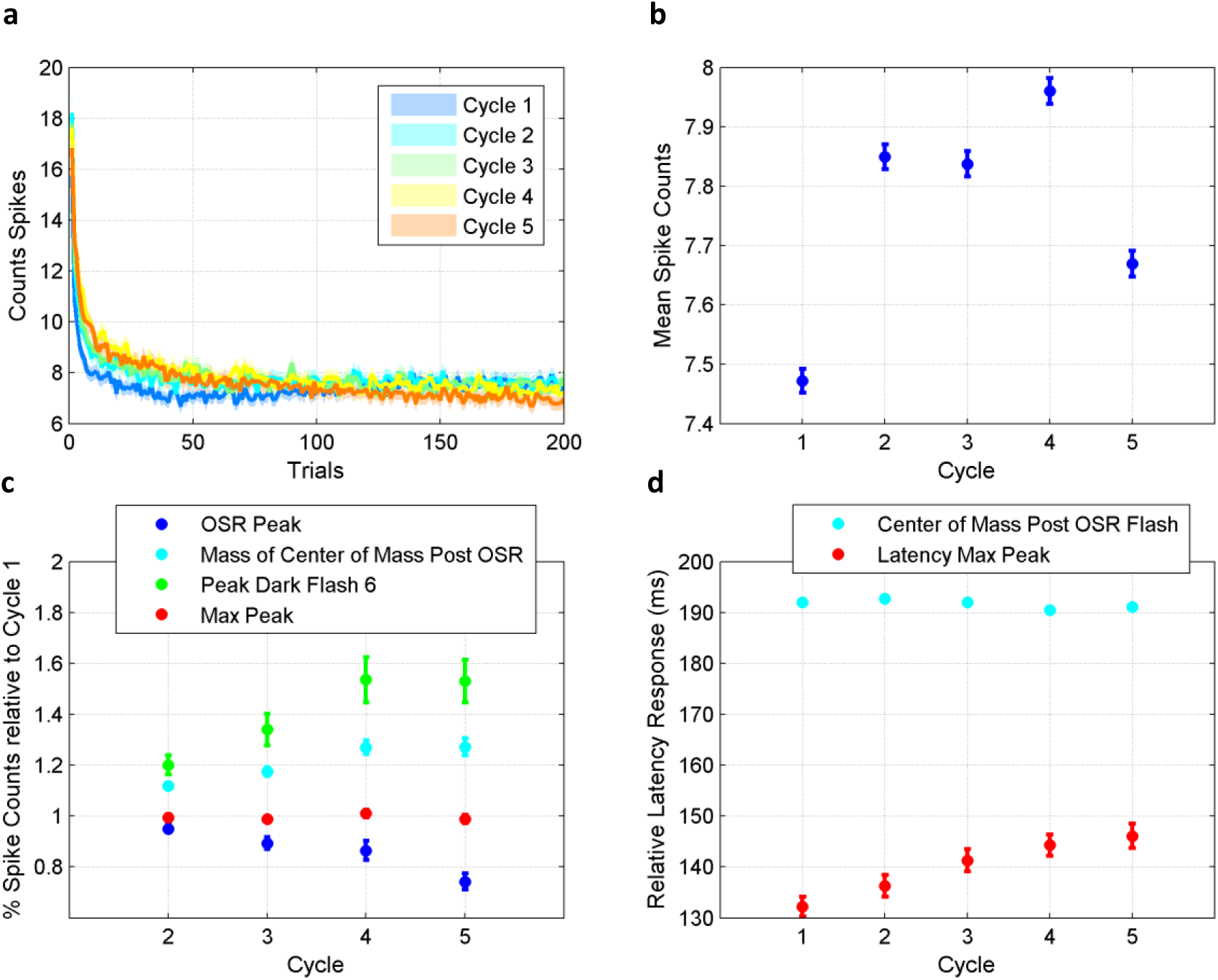
Characterization of responses cycle to cycle Habituation Protocol. Habituation protocol allows some qualitative variations on different parts of the stimulation, increasing or decreasing the neural activity. a) Adaptation Curve of the group activity per each cycle of stimulation with *τ*_1_ = 0.52, *τ*_2_ = 0.94, *τ*_3_ = 0.89, *τ*_4_ = 3.7 and *τ*_5_ = 2.66 respectively. Measures are in seconds and from an exponential fit. b) The mean firing rate evolution per each cycle of stimulation shows a tendency to increase during Habituation Protocol. Every cycle is p<0.05 (except cycle 2 and 3) with Wilcoxon test. However, the order of 0.2 for firing rate could mean something related with intrinsic variability of the system. c) Variation of firing rate response normalized for the first cycle in different parts of stimulation: OSR peak, Center of Mass for post OSR activity, Maximum peak value and peak value on flash number six (as a random reference). This plot shows how some parts of the stimulus on firing rate activity increase while other parts decrease. d) Time Latency for Maximal Peak post stimulation and Center of Mass for Activity Post Omission response. We show results for 312 units ON-OFF types from n=4 pieces of retina with Habituation scotopic Protocol. Error bar are s.e.m.

These modifications impact on the mean of the firing rate (Figure 3b). These units show differences for every cycle (p<0.05) except cycle 2 and 3, however if we look well, the order of these differences is on decimals and these variations could have relation with intrinsic variation of spike counts in retina (Berry, Warland, & Meister, 1997). These variations are apparently a decrease of the firing rate on OSR activity (Figure 3c, blue dot) that is contrasted with an increase of the activity in other parts of the pattern stimulus (Figure 3c, d), especially post OSR activity.

The OSR activity was defined as the peak on firing rate from the last flash until 150 ms (as previous works showed (Pizarro M. et al, 2014)) and post OSR activity was defined as the peak on the firing rate after 150 ms. Maximal peak is the maximal value of the firing rate post last stimulus flash independently from temporal limit. It is defined a temporal point as the center of mass for post OSR activity, and it is showed the fact that this value keeps being constant (Figure 3d) while the latency of Max peak has a tendency to increase (Figure 3d). Together, these results are suggesting a temporal shift on individual and group responses. In other words, these results are suggesting qualitative and quantitative changes on the network at temporal neuron level induced for Habituation protocol.

### Changes in Ganglion cells responses after Habituation Protocol

To see differences induced at neuron level and at population level, initial responses of ON-OFF pulses (before Habituation Protocol) and final responses (after Habituation Protocol) are compared (Tikidji-Hamburyan et al., 2015). Types of neurons are classified with two analyses: (i) How they respond to 50 trials of ON-OFF pulses (1 second per each pulse, see methods), and (ii) thanks to temporal filter clustering from STA analysis. Both classifications can be crossed and it is possible to obtain 8 different kinds of classifications. Here, 188 units for Habituation scotopic protocol (Figure S1) from three pieces of retina are reported. Then, the value of maximum peak of spike counts and time to peak (or Latency) are measured before and after Habituation protocol from simple ON-OFF pulses at the beginning and end of experiments.

With these measures, the population distribution of these units is shown for each kind of response of ON-OFF pulses (Figure 4). Again, it is possible to observe a little increase of maximal firing peak in both ON as OFF response and a bigger increase on latency ON response, after Habituation protocol. To make the comparison clearer between initial and final states, we show the paired pulse ratio (PPR) for each condition, where for “maximal firing peak”: ***Depression*** means decrease in response at final condition, Equal is all kept equal on both initial and final condition, and ***Facilitation*** is an increasing on final response. For “latency”: ***Depression*** means an increase on latency response and ***Facilitation*** is the decreasing on latency response (see methods). While maximal firing rate peak ON and OFF keeps a balance between depression and facilitation, for ON and OFF latency, it is possible to observe one or another, depression or facilitation, as dominant.

**Figure 4.**
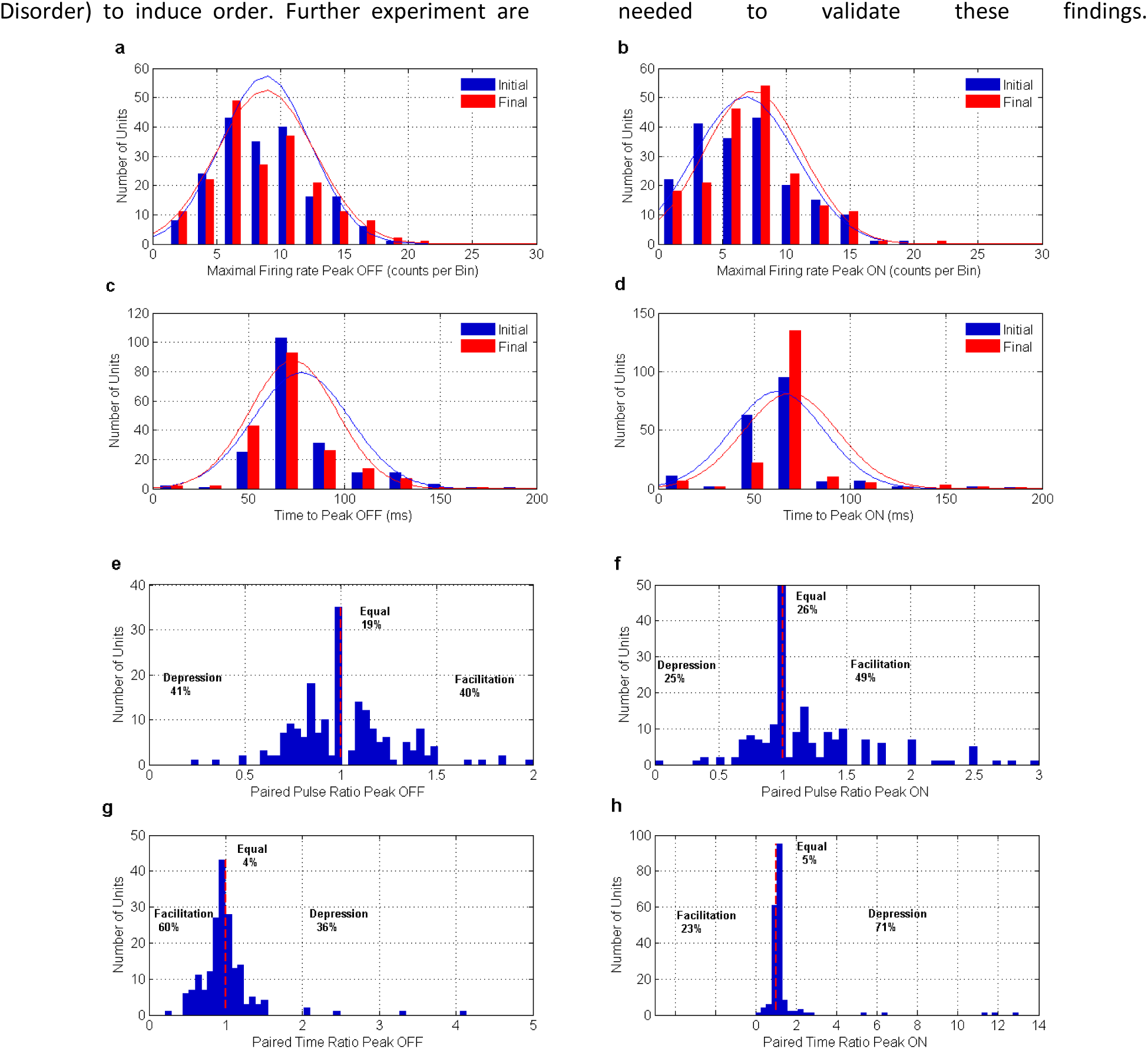
Change in the Population distribution before and after Habituation protocol. It is possible to observe changes after a habituation protocol in the population distribution of responses to a light pulse ON-OFF. a) Histogram of the maximal firing rate peak in the OFF response (p>0.05). b) Histogram of the maximal firing rate peak in the ON response (p<0.001). c) Histogram of the latency of the maximal firing rate peak in the OFF response (p<0.01). d) Histogram of the latency of the maximal firing rate peak in the ON response (p<0.001). e-f) Histogram for Paired pulse ratio Peak in OFF and ON responses respectively. g-h) Histogram for Paired Time ratio Peak in OFF and ON responses respectively. We show results for 188 units from three different pieces of retinas with Habituation Scotopic protocol. In a-b) the lines are the normal approximation distribution for each histogram. The Paired Pulse and Time Peak ratio were calculated by dividing the value in the initial state by the final value. For e-f facilitation was defined as an increase in the paired ratio (increase in spike count per bin), while for g-h facilitation was defined as a decrease in the ratio value (decrease in the latency, i.e. the final response is faster than the initial).

In conclusion, initial population distribution shows differences with respect to final distribution and effect size analyses are clearer summarizing these ideas (Figure 5 and Table 1). Figure 5a shows the mean difference of the distributions and respective confidence interval for four different protocols of Habituation. In all of these protocols, differences are very small or zero between initial and final condition on maximal firing rate peak response ON and OFF, while for latency, the effect size is considerable. Positive values mean decrease in the final condition and negative values are an increase (see methods). One interesting observation is the opposite direction for changes induced in Habituation Disorder protocol, where final ON latency decreases, unlike other protocols.

**Figure 5.**
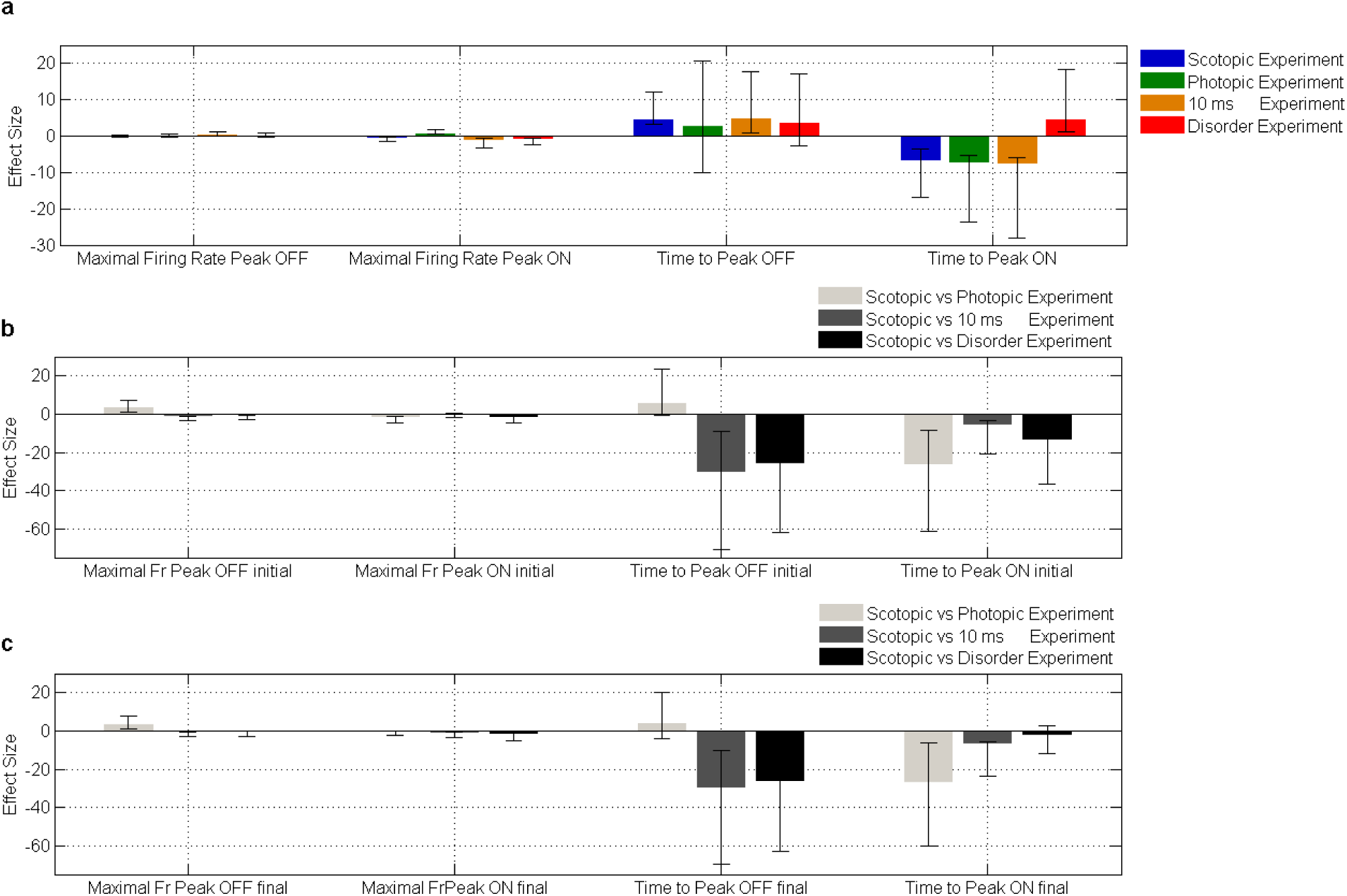
Effect Size analysis of changes on population distribution before and after Habituation Protocol by ON-OFF pulses. Effective changes are observed at different experiments, especially in the order of time response. Time to peak on OFF responses decreased for each experiment, while Time to Peak in ON responses increased on three protocols, with exception of Disorder protocol. It suggests that changes for Disorder protocol are opposite, while other three protocols affect the dynamic of the network in a similar way, but with some differences between them, induced for type of stimulus. For a correct interpretation see complementary information in Table 1. a) Effect Size between the mean population before and after Habituation Protocol. Different protocols are inducing differences of latency response: The time OFF final response decrease and the time ON final response increase (except for Disorder protocol). Analyses for four different kind of experiments: Habituation Scotopic (188 units from 3 pieces of retina); Habituation Photopic (44 units from 1 piece of retina); Habituation Scotopic 10 ms (39 units from 1 piece of retina); Habituation Scotopic Disorder (39 units from 1 piece of retina). Units in Habituation Scotopic 10 ms are the same than in Scotopic disorder experiments, and differences in direction of changes between both protocols suggest reversibility. b) Effect size comparison between Habituation scotopic protocols and other protocols at the begging of each experiment. Maximal firing rate peak for Scotopic protocol is almost the same than other protocols, but differences appear in time to peak. c) Effect size comparison between Habituation Scotopic protocols and other protocols at the end experiments. Calculation of the effect size is by the difference between the mean of the first population minus the mean of the second population. Negative values mean increased values on the second population. The bars are Confidence intervals calculated by bootstrapping technique (see methods).

Figure 5b and 5c show the comparison of effect size changes between Habituation scotopic protocol and other protocols. Figure 5b suggests that maximal spike count peak responses are similar for every experiment at initial conditions, however latency or time to peak shows differences from protocol to protocol. It suggests different initial time responses induced for each protocol. In Figure 5c, it is possible to see similar behavior than Figure 5b, with exception of scotopic and disorder protocol for time to peak ON. These findings suggest that differences induced for Habituation protocols have similar tendency for Habituation scotopic, photopic and scotopic 10 ms, while it is different for Habituation disorder, at least for ON activity.

In conclusion, repetition of one pattern at a second frequency order, as Habituation protocol, could induce changes on firing rate response, but principally on time response activity cycle to cycle. These changes are observed independently of the frequency of the pattern. Nevertheless, changes on ON latency activity are apparently driven for the pattern of stimuli (regular or irregular) and the kind of stimulation (scotopic or photopic). For example, changes induced by Habituation scotopic are more similar with scotopic 10 ms than photopic protocol, although these three protocols induce changes at population levels, mainly ON response (Figure 4 and 5), which are more similar between them than in comparison with Habituation Disorder protocol.

By this way, the phenomenon would not be associated only with repetitive patterns as OSR, if not rather with repetition itself. These preliminary results support the idea about a necessary periodic oscillation (ordered pulse pattern) between ON and OFF activity to induce disorder on retinal activity while non-periodic oscillations (Habituation

### Graph analysis for Spontaneous, OSR control and Habituation protocol activity

Topological network characterization can be applied thanks a simple pairwise correlation measures with a thresholding method based on surrogate data (see methods) (Grün, 2009; Pipa et al, 2008). Figure 6a shows an example of adjacency binary matrix for both spontaneous and stimuli driven responses, observing qualitative and quantitative differences between them. Quantitative differences can be stated using three topological network measures: GE, Q and VI.

**Figure 6.**
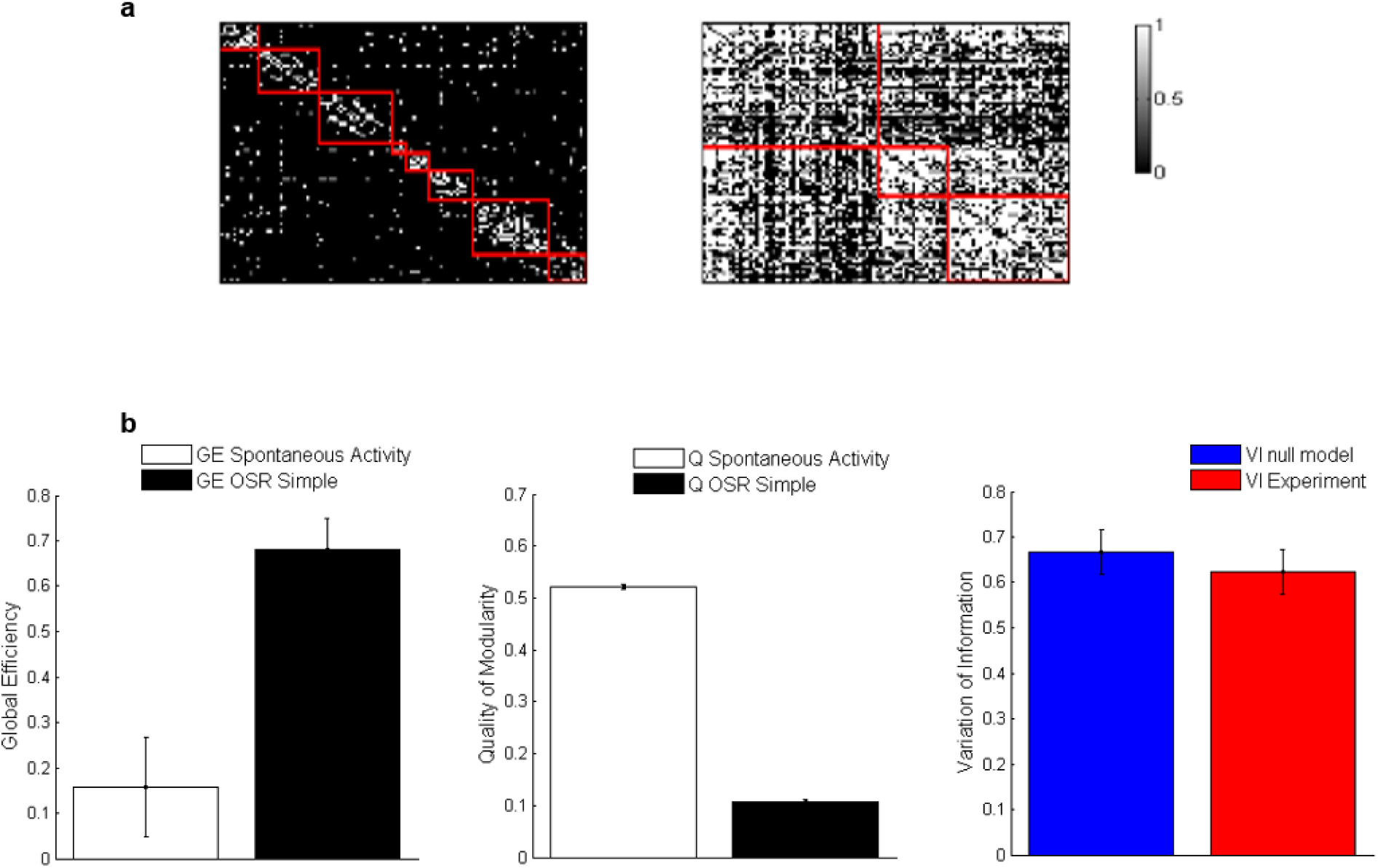
Characterization with Graph Analysis, between Spontaneous and OSR driven activity networks. **a)** Adjacency Matrix of one representative network at Spontaneous activity (left) and Stimulation activity with OSR scotopic and photopic protocol (right): 50 trials of 5 green light flash stimulation at 12.5 Hz and delay between trials of 2 second (Figure 1). Red line is representing network Modules. b) Characterization of Topological measures for Spontaneous (n=4, number of retinas) and Stimulation activity (n=5): Global Efficiency (p<0.05), Quality of Modularity (p<0.05) and Variation of the information between spontaneous and OSR activity (p=0.2 comparisons with a random network, spontaneous and OSR activity). These results show topological differences between both experimental conditions. Error bar are s.d.

GE measures are related with integration of activated patterns, or in abstract terms: some kind of information (Latora & Marchiori, 2001; Rubinov & Sporns, 2010). A value close to 1 means a bigger integration, and 0 corresponds to nothing of integration (see methods). Results shows more informative integration for stimuli OSR activity than spontaneous activity (p<0.05).

Another measure was Q (see methods), related with the degree on which networks can be subdivided into groups without overlapping nodes, these groups are called modules (Newman, 2004). The differences between both states are strong again (p<0.05) (Figure 6). This means that during OSR patterns, network activity has less potential to separate into smaller groups than in spontaneous activity. In other words, activity patterns are more segregated into spontaneous activity than stimulation activity.

Finally, VI is used to quantify difference between two network states (Meilă, 2007; Rubinov & Sporns, 2011), in this case Spontaneous and OSR activity (red bar, Figure 6b). The VI between both conditions is close to 0.7; this means a significant informative difference between partitions and modules. Comparison between spontaneous random network and OSR random network (blue bar, Figure 6b), are also computed. This random network is calculated with a rewiring process, preserving nodes and degree distribution of the original network. It is interesting to observe that there are not strong differences between VI of experimental networks and VI of random networks (p= 0.2). This last comparison was made between a random network generated from spontaneous adjacency matrix activity and another random network from OSR adjacency matrix activity. The randomization process keeps the degree distribution and nodes, while randomizes the order in connections of the adjacency matrix. One interpretation of these results is that differences are not driven by the factor randomized, i.e. changes of calculation method for random matrix can be drive significant differences, for example randomization of timestamps before adjacency matrix definition. Through this way, these findings suggest that observed properties of these networks are associated with the number of nodes and degree distribution.

On the contrary, Graph measures for Habituation protocols shows little differences cycle to cycle with any statistical different. Figures 7a and b show a representative network and qualitative evolution of its adjacency matrix between cycles, for both spontaneous and stimulation activity. GE in stimulation activity condition keeps a high value while Q value shows a tendency for increase between cycles, but it does not show any tendency during resting/spontaneous activity. However, none of these values showed a significant statistic difference cycle to cycle (p >0.05). Nevertheless, one statistical limitation is the few number of experiments (n=5) (Hentschke & Stüttgen, 2011).

**Figure 7.**
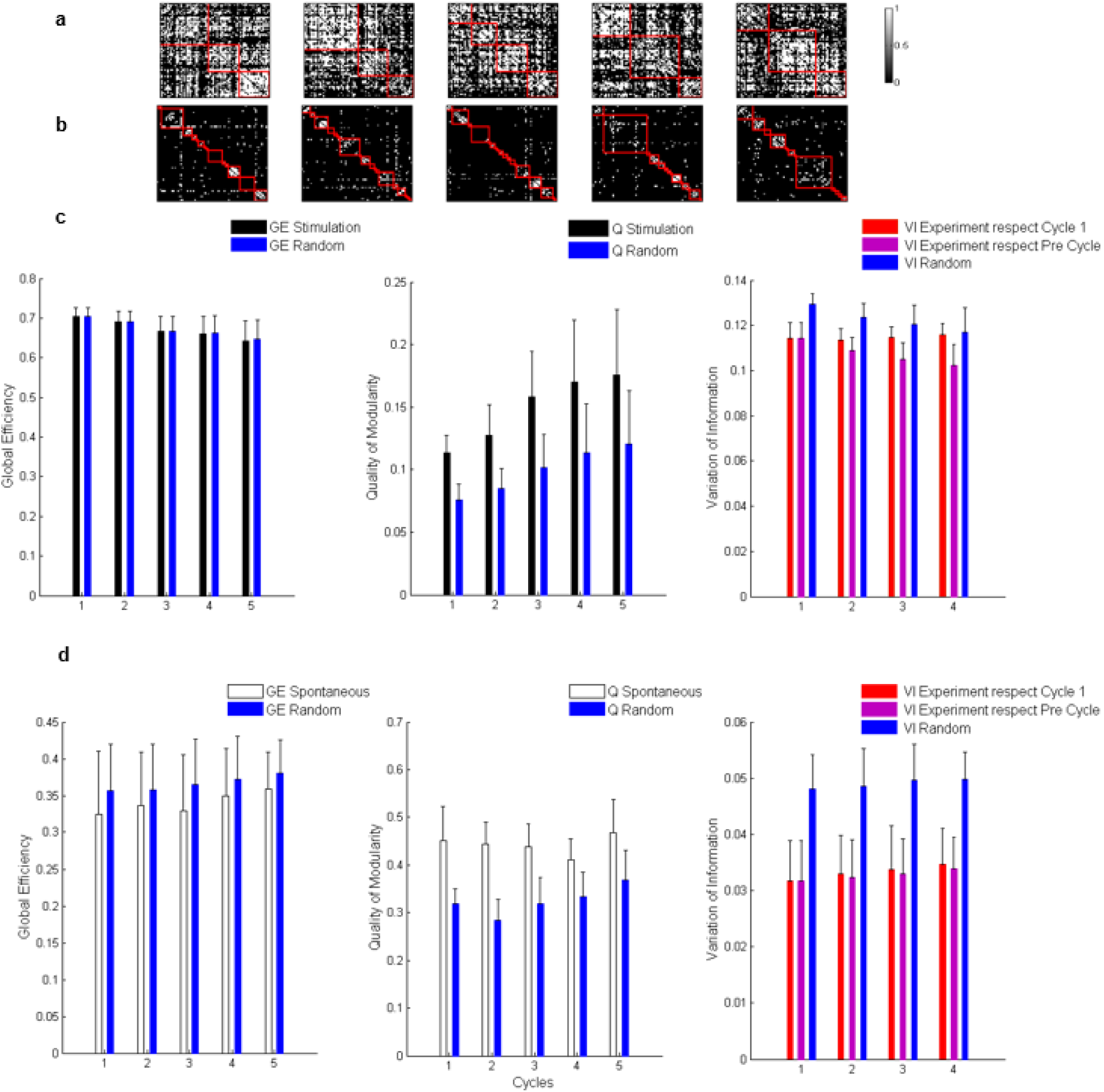
Graph characterization of Habituation Protocol. a) Adjacency Matrix of one representative network at Stimulation activity, with 74 units: 3 ON, 22 OFF and 49 ON-OFF. It is possible to observe different modules at different cycles of stimulation. b) Adjacency Matrix (same network above) in resting activity shows degenerated modular states. c) Characterization of Topological measures in Stimulation State (n=5): Global Efficiency (p>0.05), Quality of Modularity (p>0.05) and Variation of the information (p>0.2). d) Characterization of Topological measures in Resting (spontaneous) State (n=4): Global Efficiency (p>0.05), Quality of modularity (p>0.05) and Variation of the information (p>0.2). Error bar are s.e.m. In a) and b) red line is showing network Modules.

One interpretation of these preliminary results could be that “integration of information” at macro level is the same cycle to cycle, while different modular organizations are presented, i.e. the proportion of links that connect nodes between different modules are changing time to time. This can be observed as changes of Q and/or changes of number of modules (number of red square per graph, e.g. Figure 7a, b). However, these differences between modular network partitions are not impacting the global organization, as values of VI are shown. In this work is presented two different VI, i) with respect to cycle 1, ii) with respect to pre-cycle. From these VI values (Figure 7c, d), it is also possible to conclude that differences between networks partitions, due to stimuli driven activity cycle to cycle, change a little bit, in order of 10%, while for resting state activity we do not observe changes (less than 5% of differences). In other words, retinal network at resting state and its partition are globally the same at different cycles.

Thus, it is possible to define the “Robustness” of the network, when modular network partitions are more or less the same across time. Functional network robustness is defined as similar values of VI time to time and/or when variations are less than 15% [*VI_i_* ≅ *VI_f_* ∧ *VI* < 0.15] (*where VI_i_ is VI initial and VI_f_ is VI final*). One of the most interesting conclusions from these preliminary results is this robustness property in resting and stimulation driven activity.

In addition, for each cycle, changes on number of modules (see number of red square per graph in Figure 7a, and Q values which are increasing in stimulation activity) suggest different organizations of the network while they are keeping the same global topological value of GE. This is similar to degeneracy states of a physical system, where energy (in this case GE) keeps the same value for different configuration/states of the system, in this case different group organization of individual cells. This phenomenon will be called **degenerated topological states**, defined as states which preserve similar values of VI and GE across time with or without similar values of Q [*VI_i_* ≅ *VI_f_* ∧ *VI* < 0.15∧*GEi≅ GEf*∧/∨*Qi≠Qf*, or in other words, different modular organizations with the same global topological value (in this case, principally GE). A similar phenomenon on the graph context is called degenerated modular states (Rubinov & Sporns, 2011). However, in this case, Q values are constant [*VI_i_* ≅ *VI_f_* ∧ *VI* < 0.15 ∧ *Q_i_ ≅ Q_f_*]. Here, these preliminary results show degenerated modular state for resting state activity and a more general degenerated topological state for a stimulation activity in retinal network across cycles of stimulation.

These degenerated topological states and robustness could be a way how retinal networks compensate individual neural variability and keep a coherently macro response across time. These properties might be also related with healthy networks as it would present a lot of network configuration for a population response, in opposition to not healthy networks. Further experiments are needed to validate this last suggestion and compare results for retinal networks in other conditions (different stimulus or pharmacological conditions).

### Graph analysis before and after Habituation protocol

An analysis of ON-OFF pulses responses before and after Habituation protocol shows a little decreasing on GE (Figure 8b) and a little increasing on Q (Figure 8c). The VI shows a value near 0.4, this means a considerably variation between network partition between initial and final state. The effect size analysis also shows important effects induced for Habituation protocol (Figure 8e). These results suggest effective differences between both network organizations (take in count respective confidence intervals) after Habituation protocol, which are apparently compensated thanks global topological measures as GE. Figure 8 shows values for GE and Q calculated by ten times per piece of retina to avoid differences due to variability of computing algorithm (10 values per experiment, five experiments, i.e. 50 values in total). These values can impact effect size calculation, for example using standardization by normalization of the standard deviation (see methods), as it is the case in Figure 8. These are methodological differences with respect to analyses in Figure 7; however, results of VI for graph in Habituation protocol are robust. Figure 7 has small values of VI that means not considerable difference between networks (for definition VI values < 0.15) unlike results in Figure 8d.

**Figure 8.**
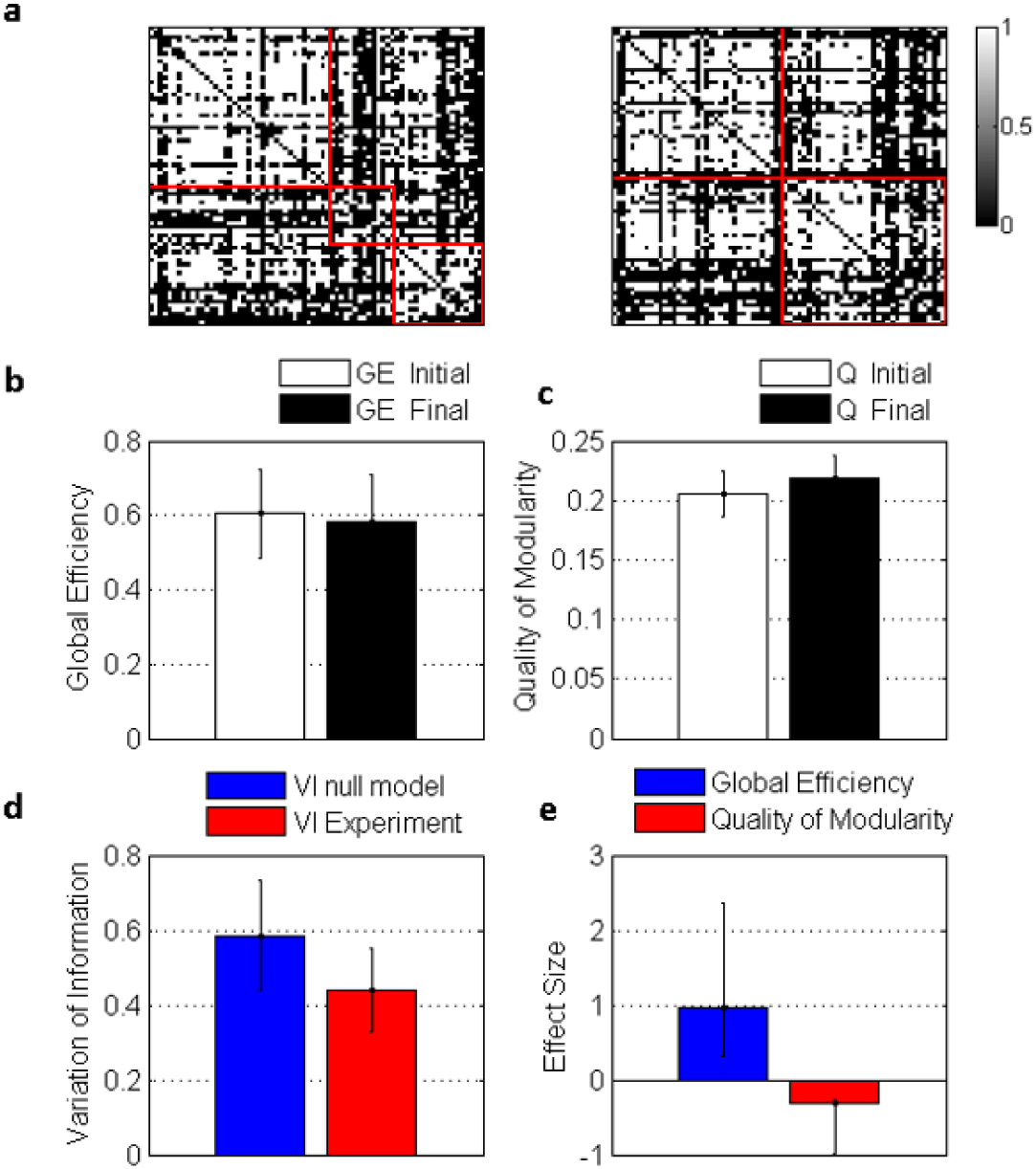
Graph Analyses for ON-OFF pulses before and after Habituation protocol. These results show similar topological graph measures in both initial and final condition. However, effective changes by effect size standardized analysis and variation of the information are also observed at modular organization level. a) Adjacency matrix at initial, before Habituation Protocol (left), and final condition, after Habituation Protocol (right), for ON-OFF pulses control protocol (example from one network with 72 units). b) Global efficiency. c) Quality of Modularity (p>0.05). d) Variation of information between initial and final condition. e) Effect Size standardized analysis. Calculation of effect size is by the difference between the mean of first population minus the mean of second population and divided by standard deviation of the difference score (differences between matching pairs of data into groups, see methods). Negative values mean increased values in the second population. The bars in e) are Confidence intervals calculated by bootstrapping technique with 1000 iterations (see methods). For a-d) bars are s.d. Values for Global efficiency, Quality of Modularity and Variation of the information were calculated 10 times per condition from Adjacency matrix to avoid potential differences due to variability of the algorithm.

## Discussion

Habituation protocols have induced effective temporal changes in both independent analyses (Figure 2, 5) and network analyses (Figure 8). Habituation protocols induced a change of latency response in independent group/population analyses and a variation of VI for graph analyses before and after Habituation protocol. Additionally, It is shown differences between spontaneous and stimuli driven activity (Figure 6), degenerated topological ability (Figure 7) and the robustness property inferred from these preliminary results.

### Variability to explain changes observed

The first option to explain changes of latency at individual and population neural level is the intrinsic variability across time of RGC activity. For example RGC have a variability of 3-5 ms when they respond to contrast or OSR protocols (Gollisch & Meister, 2008). Here, one assumption was that temporal code involve in OSR is accurate, however it is not always true for ON-OFF responses. That is why, further experiments with ON-OFF control are needed to quantify latency values and make a comparison with results exposed above. These control experiments should present pulses at different times and during same duration than Habituation protocols (approx. 40 min). Independently of this observation, the intrinsic variability cannot explain strong variability for some units (~20 ms) and it is needed a complementary explanation.

### Reversible Fatigue Process to explain changes observed

A second possibility to explain changes on latency implies loss capacity of speed information processing of external stimuli. Fatigue in RGC can be discarded, because amplitude on firing probability remains more or less constant, and it can also discard fatigue of visual pigments of photoreceptors, because even the retina is isolated from their epithelium, it is still responding to light for several hours. Another option is related with photoreceptors itself, which are involved on correlations mechanism of retina (Ala-laurila et al, 2011; Farrow et al., 2013). For example, decreasing number of photoreceptors cycle to cycle could affect response time. Nevertheless, it is possible discard this option because retinal network can recover in part original values of latency, or at least countervails their differences, as results have shown for Habituation 10 ms and Disorder experiments (Table 1, Figure 5). These experiments were performed in the same piece of retina and suggest reversibility on stimuli driven changes between Habituation 10 ms (e.g. increase latency ON) and Disorder (e.g. decrease Latency ON) (Figure 5).

Finally, the last source of changes and fatigue on retinal circuit is a possible decreasing on speed of stimulus integration of AC and/or HC, but in theory, these cells are not involved in OSR and consequently should not be involved on Habituation protocol.

### Plasticity Process to explain changes observed

Another alternative is related to plasticity process. Previous works have shown that in OSR is necessary to increase the number of flashes before the end, to increase the spike response, supporting the idea about a network training process in the order of milliseconds (Schwartz et al, 2007). As a result, some RGCs not simply respond at ended flash sequence, but for an expected flash omission, with precise time. Thus, OSR is predicting the time at which the next flash should have occurred, maintaining a short type of memory on the system that allows this prediction. One potential explanation about OSR is that a flash sequence sets up a resonance (6-20 Hz) within the retina that locks onto the stimulus frequency and it acquires a resonant frequency. When the sequence ends, internal oscillation continues and causes a large OSR (Gao et al, 2009).

If repetitive flashes as OSR are inducing a memory in some units only at milliseconds, and if these patterns are repeated trial to trial, we might be able to induce a kind of plasticity and alterations of the neural pathways and synapses, as may suggest data presented here. If this interpretation is correct, there is a possibility that these changes could be due to synaptic changes between photoreceptors and BCs synapses, BCs and GCs or both kind of synapses. However, it is not possible to know which synapses could be affected with these preliminary results.

For example, main changes are in ON latency response and OSR is associated with this way, supporting part of this interpretation. Nevertheless, as we saw differences independently of the unit, whether it has or not OSR, type ON, OFF or ON-OFF, the mechanism to explain these changes should be common for all cells types.

One option is that oscillatory stimuli induce non-normal resonance on some cells and this resonance, as a background activity, would have impact on network configuration. External oscillatory stimuli could induce oscillations on BC and interactions between BC and spike responses of RGC would modify the connectivity weight by spike-timing depending plasticity, like similar mechanism is suggested in (Masquelier et al, 2009). The intrinsic resonance properties of cells’ membrane, for example BC, would also save information as a resonator neuron memory, and this activity relative to the activity of the next layer, for example RGC, would build complex memories and fast plasticity process. In terms of resonance property, we can expect fast and reversible changes, as we see in Table 1, Figure 5.

With this last interpretation, we could also expect synaptic changes between BCs and RGCs. Typical mechanisms associated with these changes are long-term potentiation (LTP) and long-term depression (LTD) (Carew et al, 1979; Kandel et al., 2000, Caporale & Dan, 2008, Dan & Poo, 2006). However LTP and LTD are providing limited information on how the circuit and its synapses are modified by natural patterns of activity and faster learning process (Caporale & Dan, 2008; Gallistel & Balsam, 2014), (Gerstner et al, 1997; Masuda & Aihara, 2007) as it is suggested here. This is why resonance mechanism, as a kind of neural plasticity, may present an alternative to explain this missing link.

### Topological differences between Spontaneous and stimuli driven activity

Figure 6 and Figure 7 showed topological differences between spontaneous/resting and stimuli network states. Both conditions present different capacities to process information. While stimuli network states seem to have more integrated ability, spontaneous network states showed more segregation. These results are really interesting. It is known that spontaneous activity does not mean “no activity”, on the contrary, photoreceptors are liberating glutamate and some parts of this network keeps being activated, however we still found differences. These topological differences between spontaneous and stimulation activity are not apparently due to differences between scotopic and photopic spontaneous activities. Both spontaneous states are measured and results seem to be the same. It suggests that differences are due to external stimulus condition.

One interpretation could be that local correlation in spontaneous activity (in this case, local is considered some neurons near to others) (Brivanlou et al, 1998; Shlens et al, 2008; Trong & Rieke, 2008; Völgyi et al., 2013) between neighbors is stronger than non-local correlation (Mastronarde, 1989; Neuenschwander et al, 1999), while in stimulation activity, the stimuli induces a non-local correlation (Amthor et al, 2005; Simmons et al., 2013) allowing more capacity of integrate activation patterns. These results also suggest differences for other graph measures like for example clustering coefficient (whole network scale), motifs (sub-network scale), centrality (nodes scale), among others. Future works will clarify the impact of these protocols in these measures.

Additionally, a limitation to take in count is related with the definition of adjacency matrix (pairwise correlation method). On the one hand, pairwise correlation underestimates correlations value (Schneidman et al, 2006), it means that real correlation values are bigger than values of pairwise correlation. If we observe some interaction at this level, the correlation values of these interactions will increase with other correlation methods of inference. On the other hand, we know that unsupervised methods are appropriate for inference of only networks that are entirely composed of inhibitory or activating interactions but not both (Maetschke et al, 2014). If we consider, as shown above, that AC and inhibitory circuit are not involved in OSR protocols, it is possible to solve this limitation. However, different ON/OFF BC ways have inhibitory and active synapses and maybe it is not possible to avoid these interactions at all; this is why more complex inference methods based on maximal entropy or others (Gerhard et al, 2011; Wilson & Brown, 2005) should be applied to confirm these preliminary differences observed.

### Degenerate Topological States and Robustness

One important observation in Figure 7 is the degenerate topological and robustness ability: it may be related with keeping a stable or meta-stable network (Bullmore & Sporns, 2009). Results from Figure 7 and 8 showed different numbers of modules across cycles of stimulation while keeping relatively constant some topological measures. These results allow inferring robustness, degenerated topological states and coherent global response cycle to cycle. An Interesting preliminary conclusion is that retinal network might maintain informative properties thanks to the ability of generating degenerate topological states.

In other words, changes induced with Habituation, in order of minutes, are absorbed for correlation activity of the population, in a context of Graph theory. Thank to that, the network might keep coherent and robustness of network responses for external stimuli. Independently of which is the mechanism to explain these changes induced, retinal networks apparently maintains topological measures as a capacity for integrating and segregating activity patterns. It is an interesting conclusion, because, if this is correct, the retina is showing the ability to compensate individual changes in the circuit due to fatigue or plasticity across time, maintaining a coherent response. Some implications could be that in non-healthy or non-normal network this ability could be lost. This conclusion may have important implications on different dynamics among neural networks.

## Conclusion

Finally, these preliminary results have shown some interesting behavior of retinal networks when they are stimulated with the same pattern across time and how these patterns can induce changes. One suggestion for these changes is a short reversible fatigue network process, intrinsic to retinal variability. Another suggestion opens the door to search for a new oscillatory learning mechanism (Izhikevich, 1999, 2001) (Masquelier et al, 2009) (Richardson et al, 2003; Tchumatchenko & Clopath, 2014) induced by oscillatory patterns. Independently of that, in both cases, the retinal network has shown robustness on topological measures, and degenerates topological states ability across Habituation protocol.

Some perspective of this work would be complete characterization of Spontaneous versus Stimulation patterns with other topological measures, Graph theory and more complex methods of inference. Another experimental line is to relate the ability of degenerate topological state or robustness of networks, with healthy networks. In this context, the same protocol applied to non-normal network will predict a loss of robustness (i.e. changes in the Variation of information). Finally, these results should be explained by some of the existing computational models for OSR, or it would be necessary to generate new models with oscillatory plasticity and/or resonator properties, among others.

## Acknowledgment

The author appreciates valuable comments and reviews from Maria Jose Escobar, Patricio Orio and Adrian Palacios. The project was carried out thanks to Centro Interdisciplinario de Neurociencia de Valparaiso, Iniciativa Cientifica Milenio, Universidad de Valparaiso, Instituto de Sistemas Complejos, Valparaiso and CONICYT.

## Supplementary Information

**Figure S1.**
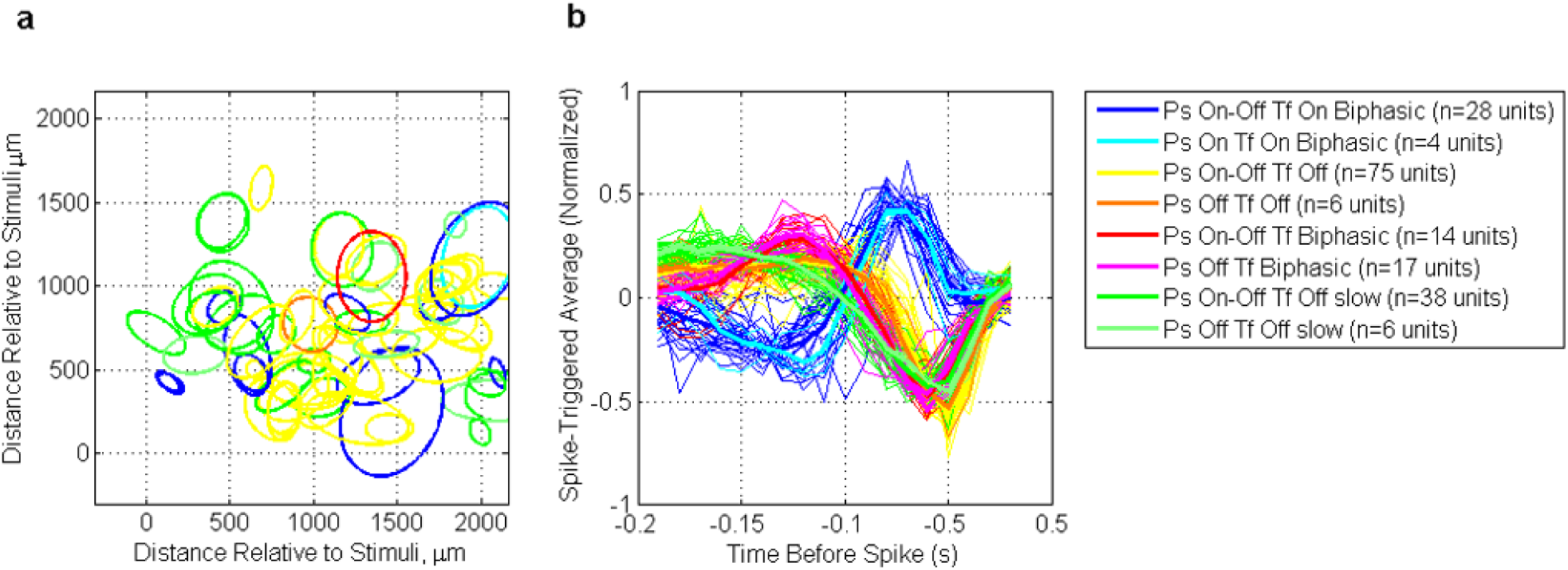
Classification of units by statistical classification criteria and STA analysis. We classified each unit in response at Pulses (Ps) of ON-OFF flashes before Habituation Protocol: ON, OFF or ON-OFF using the PSTH responses over 50 trials (see methods). ON corresponds to an onset response (between 1 ms to 1000 ms and firing rate activity>3 s.d. of the total activity); OFF corresponds to the offset response (between 1001 ms to 2000 ms and firing rate activity>3 s.d. of the total activity) and ON-OFF has both responses. For classification from the temporal filter (Tf) we used k-means to find the optimal number of clusters. In this case, we decided use just four: OFF, OFF slow, ON Biphasic, Biphasic. The final classification was merging both previous classifications. The final name is: First, the classification by the response at pulses, and second, the classification by temporal filter. a) Example of one retinal network with 72 units and their spatial filters (Receptive field). b) Temporal filter for each classification for 188 units from three different pieces of retinas with Scotopic protocol.

